# Determining and Predicting Soil Chemistry with a Point-of-Use Sensor Toolkit and Machine Learning Model

**DOI:** 10.1101/2020.10.08.331371

**Authors:** Max Grell, Giandrin Barandun, Tarek Asfour, Michael Kasimatis, Alex Collins, Jieni Wang, Firat Güder

## Abstract

Overfertilization with nitrogen fertilizers has damaged the environment and health of soil; yields are declining, while the population continues to rise. Soil is a complex, living organism which is constantly evolving, physically, chemically and biologically. Standard laboratory testing of soil to determine the levels of nitrogen (mainly NH_4_^+^ and NO_3_^−^) is infrequent as it is expensive and slow and levels of nitrogen vary on short timescales. Current testing practices, therefore, are not useful to guide fertilization. We demonstrate that Point-of-Use (PoU) measurements of NH_4_^+^, when combined with soil conductivity, pH, easily accessible weather (in this study, we simulated weather in the laboratory) and timing data (i.e. days passed since fertilization), allow instantaneous prediction of levels of NO_3_^−^ in soil with of R^2^=0.70 using a machine learning (ML) model (the use of higher-precision laboratory measurements instead of PoU measurements increase R^2^ to 0.87 for the same model). We also show that a long short-term memory recurrent neural network model can be used to predict levels of NH_4_^+^ and NO_3_^−^ up to 12 days into the future from a single measurement at day one, with R^2^_NH4+_ = 0.64 and R^2^_NO3-_ = 0.70, for unseen weather conditions. To measure NH_4_^+^ in soil at the PoU easily and inexpensively, we also developed a new sensor that uses chemically functionalized near ‘zero-cost’ paper-based electrical gas sensors. This new technology can detect the concentration of NH_4_^+^ in soil down to 3±1ppm (R^2^=0.85). Gas-phase sensing provides a robust method of sensing NH_4_^+^ due to the reduced complexity of the gas-phase sample. Our machine learning-based approach eliminates the need of using dedicated, expensive sensing instruments to determine the levels of NO_3_^−^ in soil which is difficult to measure reliably with inexpensive technologies; furthermore, crucial nitrogenous soil nutrients can be determined and predicted with enough accuracy to forecast the impact of climate on fertilization planning, and tune timing for crop requirements, reducing overfertilization while improving crop yields.

## 1. Introduction

There is a global effort to find practices for food production that can sustainably feed the population, which is expected to surpass 10 billion people by 2050.(*1*) The Haber-Bosch process enabled inexpensive nitrogen-based fertilizers to feed the booming population, with >600% increase in their use in the past 50 years.(*2*, *3*) Increased fertilization has, however, come with a great environmental cost. Approximately 12% of available arable land is now degraded, of which >240Mha (~926,000 mi^2^ or four times the area of France or state of Texas) is chemically degraded – *i.e*. contaminated with heavy metals and/or acidified, especially from nitrogen fertilizers, which interfere with nutrient mobility and uptake by plants.(*4*, *5*) Over-fertilization has visibly destroyed ecosystems by the leaching of excess NO_3_^−^ into surface waters causing eutrophication, giving rise to dead zones such as in the Gulf of Mexico.(*6*) Over-fertilization also impacts the soil microbiome.(*7*, *8*) Although this is an actively studied topic, N fertilization appears to shift relative abundance of certain microbial communities in soil, with important implication on C cycling and ecosystems(*9*).

Application of fertilizers is poorly understood and largely varied between regions and countries; for example, eight times more is applied per hectare in China than Australia.(*10*) Farmers across the globe typically rely on guidelines from their governments, fertilizer suppliers, or family know-how when deciding the economic optimum rate of fertilization to ensure maximum crop yields. Professional agronomists generally advise along guidelines and look at yields from previous years to estimate fertilizer requirements; they may also take soil samples for laboratory testing prior to sowing. Laboratory testing, however, is an expensive and slow process hence not performed regularly. Soil nitrogen (Soil-N) is crucial for high yields, and nitrogen fertilizer is the most frequently applied fertilizer. The optimal application rate is highly variable, however, since soil-N fluctuates widely with the properties of soil and weather over short timescales. Benchmark guidelines are unable to account for these variations. With the lack of data concerning the current and future nitrogen levels in soil, farmers tend towards overfertilization to protect yields, an environmentally and economically inefficient practice.(*11*–*13*)

Measurement of Soil-N is important for optimizing the use of nitrogen fertilizers and enabling spatiotemporal variable rate fertilization. Indirect spectroscopic precision farming technologies such as crop canopy sensors (*e.g*., near infrared spectroscopic cameras) can be used to approximate the N requirements of plants.(*14*–*16*) Indirect spectroscopic techniques, however, do not measure the levels of nitrogen in soil, instead they measure green light from the leaves of plants (related to nitrogenous compounds) to indirectly estimate levels of N fertilizer required. Machine learning algorithms are suitable for calibrating spectra (*e.g*., near-infrared) to soil-N.(*17*) Spectroscopic methods require plant mass (*e.g*., leaves), so the measurements cannot be performed until after germination and growth. Fertilizer, however, is usually applied just before seeds are sown, hence spectroscopic techniques rarely help in-season, and only compliment national guidelines. Using ion-selective membranes, levels of nitrogen in soil (mainly in the form of NO_3_^−^ and NH_4_^+^) can be directly detected electrochemically.(*18*) Such sensors can be integrated into Internet-of-Things (IoT) type remote sensors that can provide continuous data streams concerning levels of nitrogen in soil. To provide spatiotemporal resolution, however, many units would need to be deployed to fields.(*19*) Statistical models using machine learning are, therefore, well suited for filling in missing soil data(*20*) and forecasting them into the future.(*21*) Given each sensor node is not disposable (and expensive), they would require collection before harvest (*i.e*. labor intensive) and are susceptible to theft. They also require infrastructure investments to create a wireless network with access points etc. With the challenges such as large investment requirements, sector heterogeneity, data ownership and privacy, user acceptance and lack of interoperability, the adoption of IoT systems for soil sensing has been slow.(*22*–*24*) Ion-selective electrochemical sensors can also be produced in a small Point-of-Use (PoU) formfactor (*e.g*., Horiba LAQUAtwin, ELIT 8021). These sensors demonstrate high accuracy for NO_3_^−^ (R^2^=0.96)(*25*) and NH_4_^+^ (R^2^=0.98)(*26*) however they are delicate, relatively expensive (*i.e*. Horiba LAQUAtwin NO_3_^−^ sells for ~350 USD; each electrode ~150 USD), require sample preparation and calibration.(*27*)

In this work, we demonstrate a new and quick approach for determining crucial, but difficult to measure N-levels in soil. We combine a new type of gas-phase NH_4_^+^ sensor (to eliminate matrix effects due to the complex sample)(*28*, *29*), simulated climate data (*i.e*. rainfall and temperature), time passed since fertilization (*i.e*. number of days) and off-the-shelf soil pH and conductivity sensors with a statistical machine learning model to instantaneously and accurately determine levels of NO_3_^−^ in soil. We demonstrate that the N-levels in soil can also be predicted into the future using a long short-term memory recurrent neural network over a 12-day period. With this new approach (**Figure 1**), fertilization can be provided more precisely to improve yields, while preventing over fertilization, thus environmental degradation.

**Figure 1:**
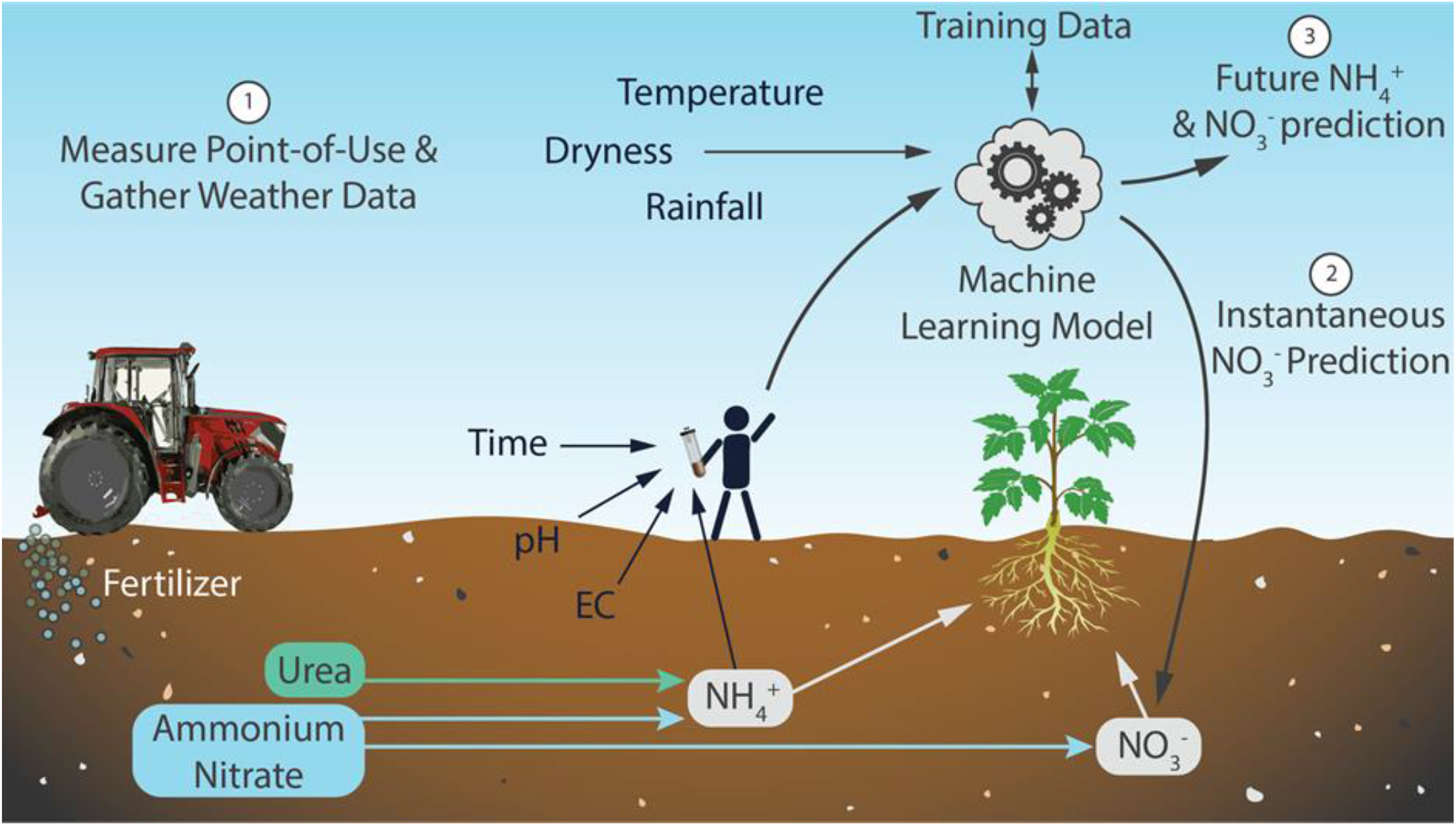
Nitrogen fertilizer is the backbone of modern agriculture, and is typically applied as urea or ammonium nitrate (we used ammonium nitrate in this work), fuelling the soil-nitrogen cycle. These fertilizers produce NH_4_^+^ and NO_3_^−^ in soil, available to be taken up by plants. We have combined point-of-use (PoU) electrical measurements (including a new soil-NH_4_^+^ sensor) with a machine learning model to quantify difficult-to-measure soil nutrients (such as NO_3_^−^) and forecast them into the future. This provides information concerning dynamics of soil nitrogen instantaneously and into the future to guide fertilization (reduce over-fertilization, understand the effect of weather, and ensure enough plant-available nitrogen to maximise crop yield) without laboratory measurements. At scale, this model could use a bare minimum of readily available input data to quantify and predict crucial outputs (*e.g*., soil macronutrients) in highly complex systems (such as soil).

## 2. Results and Discussion

### Disposable Point-of-Use NH_4_^+^ Sensor

To measure levels of NH_4_^+^ in soil, we have developed an electrical PoU sensor to accurately determine soil-NH_4_^+^ (R^2^= 0.85, limit of detection 3±1 ppm, tested up to 144 ppm) at low-cost with a large dynamic range (**Figure 2.1**). Each sensing module (*i.e*. cartridge) only consisted of a container and a disposable, chemically functionalized paper-based electrical gas sensor (chemPEGS) and 1ml 15M NaOH, costing <$0.10 (**Video SV1**)(*30*–*33*). The chemical functionalization (10 μl 0.025M H_2_SO_4_) of chemPEGS was the best compromise between precision and measurement time (**Figure S1**). Sensing of NH_3_ with chemPEGS is susceptible to interferences from other water-soluble alkaline gases, however, because NH_3_ has the highest water solubility and is the dominant water-soluble alkaline gas species in our samples due to the fertilizer (ammonium nitrate), the signal generated by the chemPEGS largely originates from NH_3_. To operate the cartridge, a soil solution was created by pressing 100 ml deionized water through 100 g of soil. A 5 ml soil solution was injected into each container. In the container, the solubilized soil-NH_4_^+^_(aq)_ is in equilibrium with solubilized NH_3(aq)_ (Equation 1), which is in equilibrium with volatilized NH_3(g)_ in the headspace of the container under Henry’s law (Equation 2).

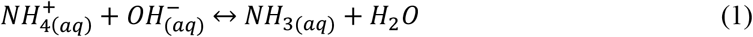

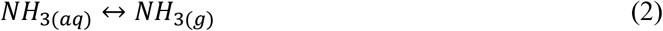

**Figure 2:**
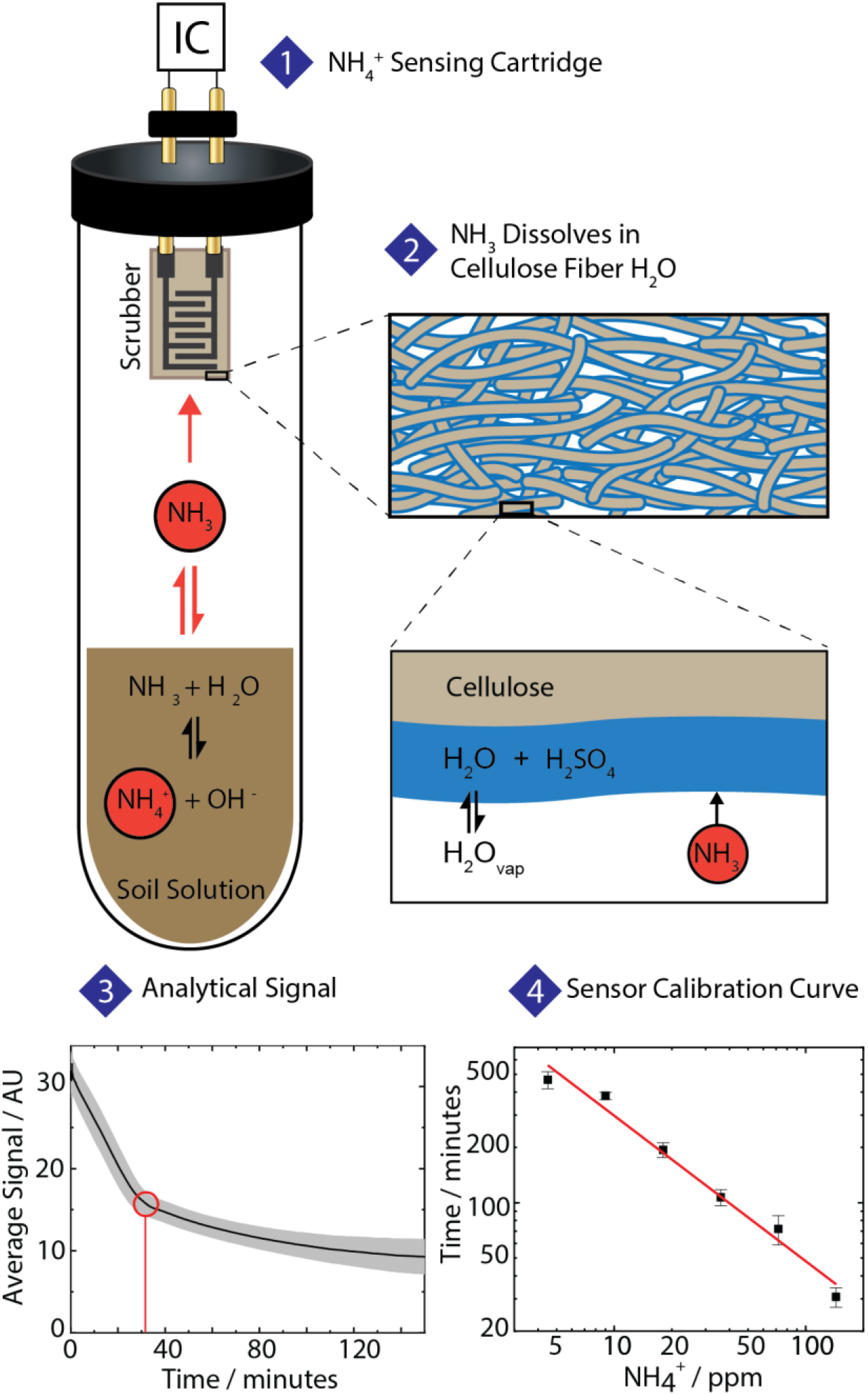
Gas-phase NH_4_^+^ sensor cartridge, consisting of a container, 1ml 15M NaOH and a disposable chemPEGS that acts as a scrubber of soil-NH_4_^+^ connected to an integrated circuit (IC) to perform impedance analysis. (**Figure 2.1**) The volatilized NH_3(g)_ dissolves in the layer of water adsorbed on the chemPEGS, (**Figure 2.2**) neutralizing the H_2_SO_4_ and increasing ionic impedance, which was measured electrically. The neutralization reaction draws out the remaining NH_4_^+^ from the soil solution to maintain the equilibrium of NH_3_ in the headspace. The time it took for neutralization to slow dramatically or complete was used as the analytical signal (see SI **Figure S3** for raw data and mathematical criteria). An example signal from 144ppm soil-NH_4_^+^ is shown in **Figure 2.3** with the analytical signal circled in red and the error shown in grey (calculated by standard deviation of n=5 measurements). We calibrated the sensor in a range of concentrations of NH_4_^+^ from 4.5-144ppm in soil fertilized with NH_4_NO_3_ (**Figure 2.4**).

The pH increased to 14 by the concentrated NaOH solution, shifts the equilibrium toward NH_3(aq)_ and ultimately NH_3(g)_. The NH_3(g)_ in the headspace of the container once again dissolves in the layer of water adsorbed on the paper sensors as described by Barandun et. al. previously,(*29*) then neutralizes the H_2_SO_4_ in paper causing an increase in the ionic impedance (mainly due to the neutralization of highly mobile H^+^ ions) of the sensor in a concentration dependent manner (**Figure 2.2**)(*28*, *34*–*36*). Neutralization of NH_3_ in chemPEGS (Equation 3) draws out more NH_4_^+^ from the soil solution to maintain equilibrium, hence the paper sensor acts as a scrubber of soil-NH_4_^+^.

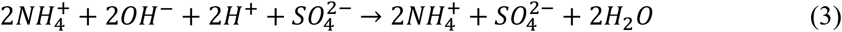

There is a decrease in ionic impedance during neutralization on the chemPEGS, which is measured electrically using our home-made electronics.(*29*) An alternating voltage (10 Hz, 4 V amplitude) was supplied across the chemPEGS, and the current passing through was measured as a voltage with a transimpedance amplifier, amplified with a gain resistor (see Supporting Information, SI, **Figure S2**). As the H_2_SO_4_ neutralization continued, the impedance of paper increased slowly and the time it took to complete or slow dramatically was used as the analytical signal shown in **Figure 2.3** (see SI **Figure S3** for raw data and mathematical criteria). Before measuring unknown concentrations, we calibrated the sensor in a range of concentrations of NH_4_^+^ from 4.5-144 ppm, in soil fertilized with NH_4_NO_3_; the calibration curve (log-log) is shown in **Figure 2.4**. Our measurements were compared to external laboratory measurements with a score of R^2^ = 0.85 (**Figure S4**). This is below reported ISE scores mentioned above, but our new sensing mechanism needs only simple and robust components for matrix free sensing and is, therefore, likely to offer more dependable results in-field. We also measured a calibration curve for NH_4_NO_3_ in water (no soil) and verified that our soil measurements are indeed from NH_4_^+^ alone (**Figure S1**).

### Time-dependent Nitrogen Dynamics in Soil

Understanding how nitrogen species evolve after fertilization, in particular the nitrification from NH_4_^+^ to NO_3_^−^, is important to growers for tailoring fertilization to climatic conditions and crop types, while reducing losses and environmental damage.(*37*) Time series data concerning dynamics of soil nitrogen were collected over short timescales (<20 days) in experiments simulating soil in a field (**Figure 3**). To reduce complexity, we did not grow any plants and investigated the nitrogen dynamics only due to microbial activity, run-off and volatilization (escape of NH_3(g)_). We placed 5.1 kg of soil (Westland Top Soil, unfertilized) in a 15 L plastic pots and stored them in the laboratory without covering their tops. In each experiment, we controlled the environmental conditions in two ways: i) adding a controlled amount of water to simulate rainfall, ii) passing an electrical current through a resistive heating wire (nichrome wire), wrapped around the containers, to control temperature uniformly. We kept the soil type (sandy loam, sieved) and amount of fertilizer added fixed for all experiments (fertilizer NH_4_NO_3_ was added in the beginning of each experiment to produce a concertation of 120 ppm, approximately equal to 241 kg/ha of NH_4_NO_3_ or 85 kg/ha nitrogen – calculation in SI). Experiments were performed for eight sets of environmental conditions spanning arid (1 mm/day rainfall) to tropical (10 mm/day rainfall) with temperatures ranging from 19-21°C (temperate) to 26-34°C (warm). Measurements of soil temperature, rainfall, pH, electrical conductivity (EC) and NH_4_^+^ were made in our laboratory (Güder Research Group - GRG). Levels of pH, EC, NH_4_^+^ were also measured in an external laboratory (NRM, Cawood Scientific), in addition to dryness and NO_3_^−^, for comparison and training of the machine learning model. Although when building the machine learning model we relied on the dryness values provided by the commercial laboratory, dryness is highly correlated with rainfall and temperature (in our dataset, temperature and rainfall predict dryness using linear regression with R^2^ = 0.86, see **Figure S5** and the equation in the SI) hence can be estimated using these two metrics without needing further analytical measurements.

**Figure 3:**
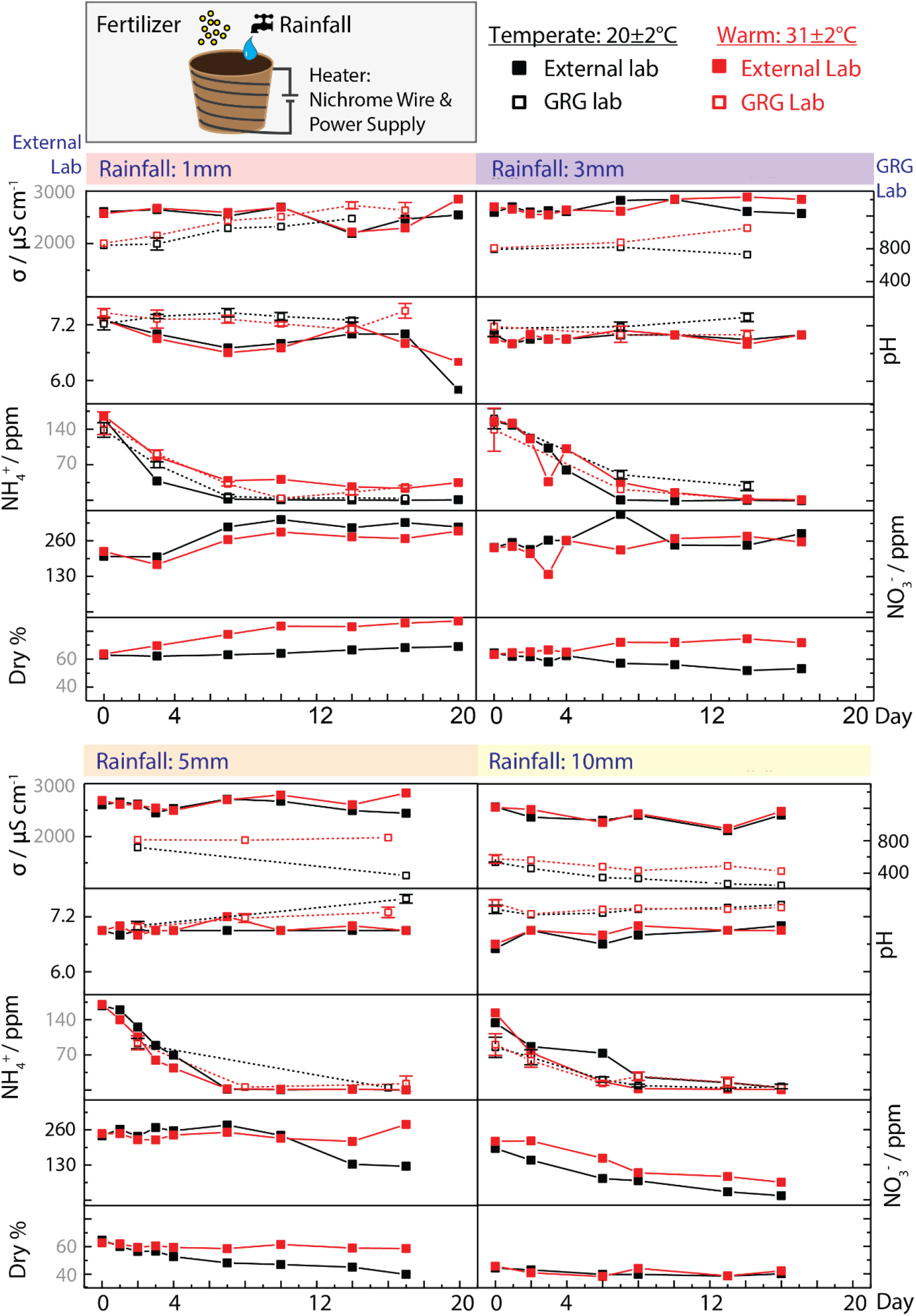
Time series data of soil nitrogen dynamics were measured over short timescales (<20 days), where each corresponds to soil under different environmental conditions. We controlled rainfall and temperature by adding a controlled amount of water and passing current through a resistive heating wire (top of **Figure 3**). Environmental conditions span arid (1mm rain/day) to tropical (10mm rain/day), temperate (20±2 °C) and warm (31±2°C). Initial fertilization with NH_4_NO_3_ was fixed at 120 ppm. Measurements of soil temperature (n=3), rainfall, pH (n=5), EC (n=5) and NH_4_^+^ (n=5) were made in our laboratory (GRG) with errors corresponding to the standard deviation, and pH, EC, dry %, NH_4_^+^ and NO_3_^−^ were measured in an external laboratory (NRM) for comparison and training the machine learn model.

#### Dynamics of soil NH_4_^+^

In all time dependent soil experiments, the level of NH_4_^+^ dropped rapidly over time, levelling out after about a week, independent of the environmental conditions. Temperature played a considerable role only in the case of 1 mm/day rainfall/ in which the NH_4_^+^ levels settled at ~50 ppm for warm conditions, in comparison to ~0 ppm for temperate conditions. In all other scenarios, temperature or rainfall only slightly affected the NH_4_^+^ dynamics without large differences in the trends. Decreasing levels of NH_4_^+^ result from multiple processes, such as nitrification (*i.e*. conversion of NH_4_^+^ → NO_2_^−^ → NO_3_^−^) or environmental losses (leaching or volatilization), that run in parallel; however, the extent of each process might vary with environmental and soil conditions. Soil dehydration tends to limit nitrification, by restricting substrate supply to microbes and lowering activity of enzymes,(*38*) which may explain retention of NH_4_^+^ at higher temperatures (and low rainfall). This observation is further supported by the fact that the levels of NO_3_^−^ were lower for warm conditions than temperate conditions.

#### Dynamics of soil NO_3_^−^

Nitrification is a complex, aerobic microbial process affected by temperature, moisture, levels of O_2_, pH and of course availability of NH_4_^+^ among other things (*e.g*., nitrifier populations).(*37*) We observed that, while at 1 mm/day rainfall the level of NO_3_^−^ increased compared to the initial (day zero) concentration, for 3 mm/day it remained relatively unchanged both for warm and temperate conditions. For 5 mm/day rainfall in warm conditions, the levels of NO_3_^−^ only slightly increased toward the end of the experiment. For temperate conditions the concentration of NO_3_^−^ nearly halved with a rapid drop after day 10. For heavy rainfalls (10mm/day), the concentrations of NO_3_^−^ dropped toward zero in a linear manner over the course of the experiments. From these experiments, it could be concluded that the optimum point for maximum nitrification and retention of NO_3_^−^ in soil occurs in temperate and drier conditions, which are consistently more favourable than warm and wetter conditions. The reasons behind these trends may differ, however, depending on the conditions. While the run-off caused by the heavy rainfall (*i.e*. 10 mm/day) may physically leach NO_3_^−^ away (the excess water was pouring out from the bottom of the pots), less rainfall (5, 3 mm/day) may hinder penetration of O_2_ into the soil (*i.e*. waterlogged soil) therefore reduce nitrification, especially if the climate is temperate so that not enough water is removed from the soil to allow oxygenation.(*39*) The optimal temperatures for nitrification are typically reported between 24-27 °C,(*40*) in line with our observations. In the experiments where the dryness of soil did not increase, temperature did not have a large effect, evidenced by the first 4 days of the experiment with 3 and 5 mm/day rainfall. Dryness (*i.e*. rainfall + temperature), therefore appears to be a more important factor in determining the levels of NO_3_^−^ than temperature alone.

#### Dynamics of soil EC and pH

EC and pH were measured to investigate their correlation with soil nitrogen under different environmental conditions. Due to technical difficulties, we were unable to complete the EC and pH measurements for all samples in a single day, hence missed measurements which were to be performed in our laboratories. Nevertheless we did not observe any major trends in pH or EC regardless of rainfall or temperature except for the experiments with 1 and 10 mm/day rainfall. For 1 mm/day rainfall, the EC only slightly increased and pH slightly decreased over time. Ammonium based fertilizers are known to acidify soil therefore decrease pH.(*41*, *42*) With an increase in the concentration of mobile NO_3_^−^ ions in soil, EC is also known to increase.(*42*) When the rainfall was increased to 10 mm/day, however, the run-off leached out ionic species from the soil, in turn reducing EC of soil without affecting pH. The EC and pH measurements performed in our laboratory and externally did not correlate to the degree we expected, although the instruments used in our laboratory were calibrated weekly with calibration solutions to produce reliable measurements. Upon investigation, we found out that the difference in sample preparation was the likely culprit behind differences in the results. The external laboratory dried the soil samples before taking a fixed weight and mixing with water for measurements, whereas we took the samples directly from the pots without drying and mixed with water, which caused varied values for EC and pH. In any case, in the context of this work, these differences in sample preparation did not affect the underlying trends in the data generated by the external laboratory and such small errors may happen under real experimental conditions at the point-of-use (hence the entire system should be robust enough to absorb these errors).

### Prediction of levels of nitrogen in soil using machine learning

Retention, conversion or loss of nutrients added to soil is a complex function of rainfall, temperature, pH, microbe populations, soil type etc. This complexity renders creation of deterministic models to understand the relationship between nitrogenous species and their levels in soil, difficult (if not impossible) after some time, even if initial concentrations are known. We have, therefore, attempted to create a statistical model using (existing) machine learning (ML) approaches to predict levels of hard-to-measure NO_3_^−^ in soil using information concerning weather (*i.e*. rainfall and temperature), time since fertilization, pH, EC, and NH_4_^+^ (**Video SV2**).

Using supervised ML, we attempted to predict the level of NO_3_^−^ in soil instantaneously, and both NH_4_^+^ and NO_3_^−^ into the future (see **Figure S6** for the ML prediction process flow). We used the measurements performed by the external laboratory (**Figure 3**) as a training set (data processing for ML described in SI). The performance of the model was then tested either with data from the external lab or data generated by the PoU sensors in our lab as inputs. Training data matching the same environmental conditions (temperature and rainfall) as the test inputs were removed, so the model was always tested on unseen environmental conditions. Features were ranked in order of importance (by XGBoost, **Figure 4.1** top left), where soil dryness, time since fertilization and NH_4_^+^ were the most important. We compared combinations of features, regressors and tuning parameters exhaustively (by grid search) to find the best general regressor to estimate the levels NO_3_^−^ instantaneously (see **Figure S7** for R^2^ scores for each set of environmental conditions). We have determined that the K-nearest-neighbors (Knn) algorithm, trained on all 7 features with tuning parameters of k=14, leaf size=1 and p=1, can predict instantaneous levels of NO_3_^−^ with R^2^ = 0.63 using external lab results for training and our lab PoU sensors for test input (**Figure 4.1** bottom left). Using the same model, but with external lab results as test inputs, removes the impact of inaccuracy from our lab (PoU) sensors, resulting in R^2^=0.68 (**Figure S8**). We have also determined that XGBoost regressor produced the best predictions for dryer soils and Knn for wetter. Taking our lab PoU sensors as test inputs, and the optimal regressor and tuning for each set of environmental condition offers even better performance, giving an optimised score of R^2^_av_= 0.70 (score averaged across each set of environmental conditions – **Figure 4.1** bottom left). The same process, but with external lab result as test inputs, produces on optimized best-case scenario score of R^2^_av_= 0.87 (**Figure 4.1** bottom right). This score for predicting levels of NO_3_^−^ in soil is comparable to direct measurements using optical (R^2^ = 0.83; Fourier-transform infrared spectroscopy) or electrochemical methods (R^2^ = 0.96; ion selective electrodes) as reported in the literature.(*26*, *43*) This result was pleasantly surprising given that no additional hardware was required for determining levels of NO_3_^−^ with high accuracy. We have also tuned a Knn model (k=11, leaf size=3, p=1) to predict levels of NO_3_^−^ in soil using only the most basic inputs – days since fertilization, rainfall and temperature (*i.e*. requiring no soil sensors at all) which yielded R^2^ = 0.54 (**Figure S9**).

**Figure 4:**
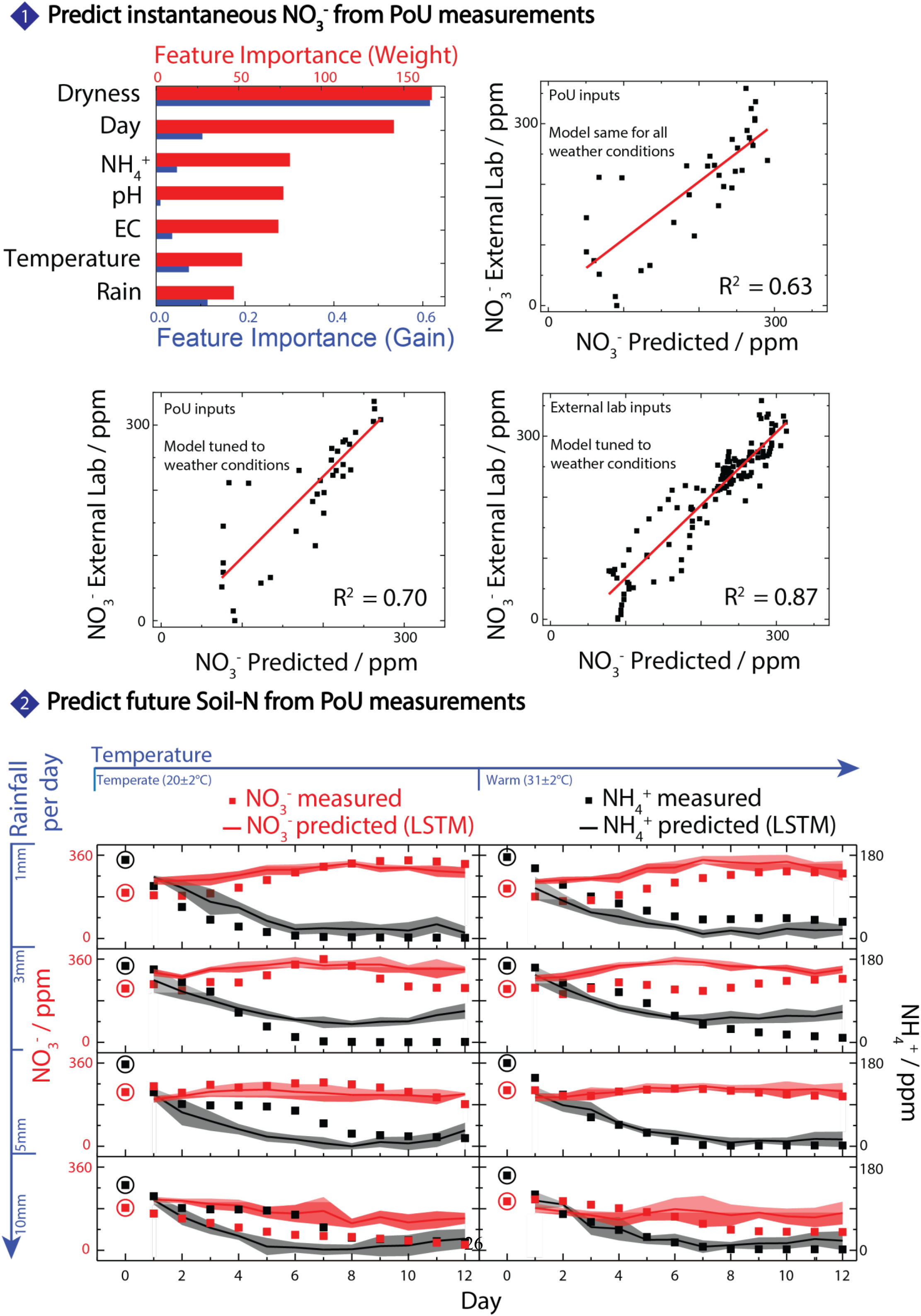
Instantaneous Soil-NO_3_^−^ was predicted with instance based and ensemble learning regressors. Features were first ranked by importance to the XGBoost model, calculated by weight (number of times the feature occurs in the trees) and gain (each feature’s contribution to each tree), shown in **Figure 4.1** (top left). The best performing regressor for all environmental conditions is Knn, which predicts NO_3_^−^ using only PoU sensors from our lab as test inputs with R^2^=0.63 (**Figure 4.1** top right). Optimizing the model for each set of environmental conditions (regressor and tuning for each shown in **Figure S7**) improves the score to R^2^=0.70 (**Figure 4.1** bottom left). Using external lab data as test inputs (removing any inaccuracy from our PoU sensors) gave optimized predictions for NO_3_^−^ with R^2^=0.87 (**Figure 4.1** bottom right). The timeseries dataset was used to train a long short-term memory recurrent neural network (LSTM-RNN) model to forecast NH_4_^+^ and NO_3_^−^ into the future, also for unseen environmental conditions. Models were retrained for each desired forecast time (1-12 days into the future) and comparing predicted to real values over the 12 day period gives a score of R^2^_NH4+_ = 0.64 and R^2^_NO3-_ = 0.70 (**Figure 4.2**).

Although determining the concentration of NH_4_^+^ and NO_3_^−^ in soil at any given moment is important (as described above), from an operational point of view, it would also be useful to know what the levels of soil nitrogen (*i.e*. NH_4_^+^ and NO_3_^−^) will be in the future from a single measurement to create a precise schedule for future fertilization. Soil, however, introduces a memory effect: nutrient levels today depend on the nutrient levels and other factors from yesterday (property X at time t will be a function of X at t-1). Forecasting of soil-N into the future must, therefore, consider time and sequence of data, and possess a degree of memory, for multiple correlated features. Using the time-series dataset generated by the external lab, we trained a long short-term memory recurrent neural network (LSTM) model (another supervised ML algorithm) to forecast NH_4_^+^ and NO_3_^−^ into the future for unseen environmental conditions. We tuned the model using grid search, minimizing root-mean-squared error using time lag and model hyperparameters (training epochs, batch size, number of neurons). The optimal tuning was time lag=1, epochs=50, batch size=3 and number of neurons=3. The dataset was first concatenated into one multivariate time series. Each time series was then removed sequentially, and the model trained to predict the removed time series from the remaining data. Models were retrained for each desired forecast time (1-12 days into the future). Predictions for longer time periods were distorted by subsequent time series in the concatenation. Comparing predicted to real values over the 12-day period gives a score of R^2^_NH4+_ = 0.64 and R^2^_NO3-_ = 0.70 using only the initial concentrations for NH_4_^+^ and NO_3_^−^ on Day 0 which demonstrates efficacy even with our limited training dataset (**Figure 4.2**, with R^2^ plots in **Figure S10**). In essence, by measuring NH_4_^+^, EC and pH in the field and gathering other environmental data from public sources, levels of NO_3_^−^ can be estimated for today and both levels of NH_4_^+^ and NO_3_^−^ into the future.

## 3. Conclusions

In this study, we demonstrated that it is possible to estimate the levels of hard-to-measure chemicals in soil using easily accessible soil/climate data and ML models. This entirely new strategy allows determining and predicting levels of nitrogen (NH_4_^+^ and NO_3_^−^) in soil, both instantaneously and into the future. We have produced the first soil nitrification dataset that provides enough temporal resolution (≈3 day measurement frequency), for a range of conditions, to train a ML model. The strength of our approach is that we primarily use, inexpensive/easily accessible tools for the soil measurements (pH and EC meter with the exception of a new PoU paper-based, gas-phase NH_4_^+^ sensor developed in this work) and publicly available weather data (rainfall and temperature; in this study we simulated weather in a controlled manner) to estimate the levels of soil nitrogen through ML. The method presented is remarkably high performance such that concentration of instantaneous soil-NO_3_^−^ can be estimated using PoU inputs with R^2^_av_=0.70, and external laboratory inputs with R^2^_av_=0.87 (comparable to existing high performance NO_3_^−^ sensors) without the need for additional hardware. Using a LSTM model, the levels of NH_4_^+^ and NO_3_^−^ can also be forecasted 12 days into the future, for unseen environmental conditions, with R^2^_NH4+_ = 0.64 and R^2^_NO3-_ = 0.70. Furthermore, the paper-based, disposable, gas-phase NH_4_^+^ sensors (*i.e*. chemPEGS) developed in this work could also be used alone at the PoU without the ML model or other sensors if instantaneous detection of NH_4_^+^ is needed alone. The approach presented in this work has the following three weaknesses:

i. The supervised ML algorithms used for the prediction of soil-N require a training dataset, meaning prior measurements/climate data are needed to make the estimation algorithms work. This problem could partially be resolved by using data for soil nitrogen already published in the literature to create a training dataset. A training dataset could also be created using the PoU sensor toolkit described in this work in addition to occasional measurements of soil-NO_3_^−^ in an external laboratory. We expect that performance of the algorithms will increase over time as more data are generated using the sensors and laboratory measurements. Although leave-one-out cross validation was used for training, the lack of a validation dataset may have resulted in overfitting of the hyperparameters to the weather conditions in **Figure 4.1** (bottom left and bottom right). The LSTM approach also concatenated all training data into one long multivariate time series, resulting in a model that would only predict cyclical patterns if allowed to predict longer times than the length of the input time series (t ≥ 16 days). This problem may be addressed by treating each set of environmental conditions as panel data (with separate multivariate time series for PoU measurements in different locations/environmental conditions), linking between panels and training over longer time periods.
ii. chemPEGS (for measuring NH_4_^+^ at the PoU) is expected to be cross-sensitive to other alkaline gases and currently takes a long time to perform a measurement; 30-450 minutes for 144-4.5 ppm NH_4_^+^. chemPEGS, however, demonstrated sufficient performance for measuring soil-NH_4_^+^ as it is most sensitive to NH_3(g)_ due to its high water solubility. The time it takes to produce a result could also be reduced by measuring the rate of change of the impedance during neutralization or training a predictive machine learning model on short measurement times. The sensitivity of chemPEGS could also be improved by using a lower concentration of H_2_SO_4_.(*29*, *30*, *44*)
iii. The dataset generated in this work is limited (sparse) and does not include various scenarios such as sudden changes in weather, different types of soils and fertilizers (*e.g*., urea). The current work also does not include crops, which would draw nitrogen from the soil and affect nitrogen dynamics. A limited dataset may explain the unusual optimized LSTM settings for batch and number of neurons (both 3) Further work is needed to create a model to predict nitrogen uptake by plants.

The impact of this work is that growers can instantly determine crucial soil nutrients using only point-of-use measurements and weather data, and forecast nutrients into the future to build better fertilization plans. This would ensure that appropriate nutrients are present, when needed, by the crops. This approach could enable precision farming of a new caliber (with significantly lowered capital investment), reducing fertilizer requirements, soil degradation and eutrophication, while improving crop yields. Furthermore, we hope this approach will extend to complex media other than soil, where simple chemical measurements and easily accessible data, combined with machine learning, can be used to predict, and forecast crucial outputs in healthcare, food and environmental monitoring.

## 4. Experimental Details

### Soil experiments

Top soil with sandy loam texture (69% sand 2.00-0.063mm diameter, 25% silt 0.063-0.002mm diameter, 6% clay <0.002mm diameter, density 774 g/l measured in NRM Laboratories, part of Cawood Scientific, United Kingdom) was purchased from Westland and used in the experiments without further modifications. For the soil experiments performed in our laboratory, the water-soluble compounds and small particles were extracted from the soil samples by mixing 100ml diH_2_O with 100g of soil, and pressing with a potato press (VonShef). The solution extracted was used in the subsequent, pH, EC and NH_4_^+^ measurements in our laboratory. Soil samples (200 g), for the measurements at the external laboratory (NRM), were extracted from the soil pots and stored in a Ziploc bag (placed inside a cool box along with cooling element) which were collected and analysed within 2 days. Different to our method of handling, the external lab used a soil-to-water ratio of 1:2.5 as they dried the samples before processing to improve consistency (we did not do this, which caused issues surrounding unmatching results between the external measurements and measurements performed by our group). Levels of soil nitrogen were measured colorimetrically by the external laboratory. NH_4_^+^ was reacted with alkaline hypochlorite and phenol to form indophenol blue. Sodium nitroprusside acted as a catalyst in formation of indophenol blue which was measured at 640nm. NO_3_^−^ was reduced to nitrate using cadmium in an open tubular cadmium reactor. A diazo compound formed between nitrite and sulphanilamide, which was coupled with N-(1-Napthyl)ethylenediamine dihydrochloride to give a red azo dye, measured at 540nm. For all soil experiments, soil was weighed into pots of 5.1 kg, and fertilized with 51ml 0.665M (12,000ppm) NH_4_NO_3_ while mixing thoroughly, resulting in soil at approximately 120ppm NH_4_NO_3_.

### Fabrication of chemPEGS

chemPEGS with carbon electrodes (No. C2130925D1 conductive carbon ink, 55/45 wt % with No. S60118D3 diluent from GWENT Group) were screen-printed on chromatography paper (WhatmanTM, grade 1 chromatography paper, 20 cm × 20 cm, 0.18 mm thickness) and dried overnight at room temperature to remove excess organic solvents from the electrodes. The design of the electrodes consists of three interdigitated electrodes with 1 mm spacing between each finger which was optimized to increase sensitivity – *i.e*. we would like to keep the sensors as resistive as possible up to the level where our electronics could handle the low-currents. To contain the H_2_SO_4_ added in a certain region within paper, wax was deposited around the electrode area to provide a hydrophobic barrier. We printed wax designs using a Xerox ColorQube 8580 printer on to Office Depot transparent acetate sheets, then heat-transferred to the chemPEGS substrate with a Vevor HP230B heat press (180 °C). 10μl 0.025M H_2_SO_4_ was then drop casted on the chemPEGS before use, to neutralize the ammonia gas. An array of batch processed chemPEGS sensors is shown in **Figure S11**.

### Gas-phase measurement of NH_4_^+^ with chemPEGS

A gold plated card-reader (Midland Ross CD6734 BRN 34 Way IDC Ribbon Cable Card edge Connector) was inserted into a screw cap of a 50ml centrifuge tube (VWR) and sealed with a glue gun. Using the electrical contacts on the back (left outside the tube), the card-reader was connected to a custom-built printed circuit board containing electronics that can apply a10Hz, 4V_p-p_ signal across the chemPEGS. chemPEGS was inserted into the card-reader and placed in a sealed centrifuge tube containing 1ml 15M NaOH. The current produced as a result of the voltage applied was converted to a voltage again by an operational amplifier-based transimpedance amplifier (with a ratio defined by a gain resistor); the voltage signal was subsequently recorded by an Arduino Due using its onboard Analog-to-Digital Converter and communicated to a near-by PC over a serial link. Once the electrical signal (current passing through the sensor) stabilized (*i.e*. the paper substrate reached an equilibrium with the humidity inside the tube), a 5ml soil solution was injected into the tube using a syringe and changes in the electrical current was recorded as the analytical signal.

### Calibration of chemPEGS for measuring soil-NH_4_^+^

The increase in impedance (calculated using Ohm’s Law) of the chemPEGS was measured over time during neutralization of H_2_SO_4_ that was drop casted previously. Calibration was first performed without soil for a range of NH_4_NO_3_ concentrations (4.5, 9, 18, 36, 72, 144 and 287ppm), and a range of H_2_SO_4_ concentrations (0.1, 0.05 and 0.025M), shown in **Figure S1**. Calibration was attempted by comparing concentration of NH_4_NO_3_ to the total change in the impedance of chemPEGS, and the time taken for impedance to stop increasing or slow dramatically (**Figure S3**). A concentration of 0.025M for H_2_SO_4_ gave the fastest and most precise measurements (**Figure S1**). Calibration in soil solution was performed for a range of concentrations (4.5, 9, 18, 36, 72, and 144ppm) of NH_4_NO_3_ spiked in the soil sample and then extracted. A calibration curve was fitted as NH_4_^+^_[ppm]_ = (Time_[minutes]_ × 5.43e-4)^−1.26^ with R^2^_Cal._ = 0.96 (**Figure 2.4**).

### Control of rainfall and temperature

Rainfall was fixed at 1, 3, 5, or 10 mm/day, implemented by adding a daily equivalent (pots were watered every 2 days) of 57ml, 172ml, 286ml and 573ml respectively to a pot area of 573cm^2^. Temperature was controlled by wrapping pots containing soil with nichrome wire (purchased from Amazon) and applying a 36V potential, resulting in an electrical current of 1.5A supplied from two Tenma 72-8350A power supplies in series. Soil temperature was measured at 3 points (centre, edge and in between) and averaged to estimate the temperature of soil periodically, using a Silverline 469539 Pocket Digital Probe Thermometer.

### Measurement of EC and pH of soil

Using a Hanna Instruments HI5222-type benchtop EC/pH meter, the pH and EC of the solution extracted from the samples of soil were measured. Each sample was measured five times and the readings were averaged to reduce error.

### Machine learning model

All computational work was performed using Python (3.6) in PyCharm integrated development environment. For modelling and optimization, we used the following core packages: Keras API for Tensorflow (LSTM model), Scikit-learn (ensemble and Knn regressors), XGBoost, pandas and NumPy.

## 5. Acknowledgements

We would like to thank EPSRC (EP/R010242/1), Innovate UK (33486), Cytiva (formerly General Electric Healthcare Life Sciences) and Imperial College, Department of Bioengineering for their generous support. Firat Güder and Max Grell thank Imperial College Centre for Processable Electronics and Centre for Doctoral Training in Plastic Electronics, Professor Sophia Yaliraki from Imperial College Department of Chemistry, Professor Thomas Bell from Imperial College Department of Life Sciences, and Dr Tom Schaul from DeepMind Technologies Ltd. Firat Güder also acknowledges Agri Futures Lab. Michael Kasimatis acknowledges EPSRC DTP (1846144). Alex Silva Pinto Collins acknowledges BBSRC DTP (2177734).

