## Supporting Information for "Determining and Predicting Soil Chemistry with a Point-of-Use Sensor Toolkit and Machine Learning Model"

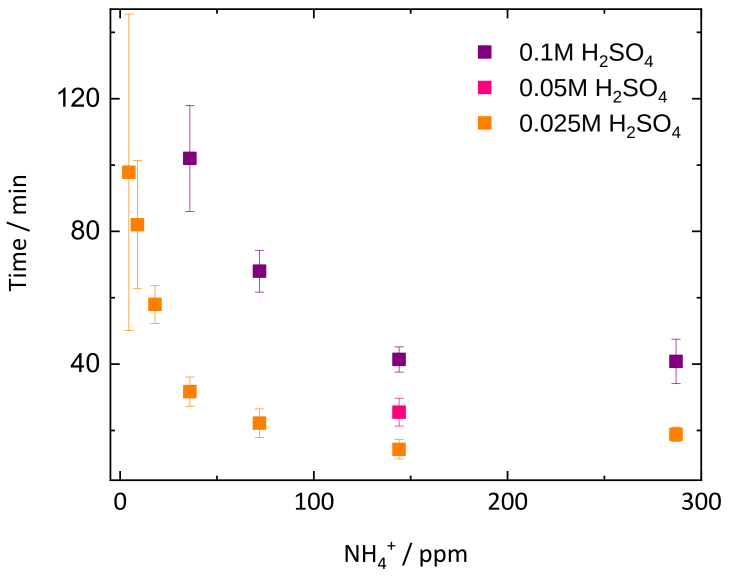


**Figure S1**

We measured calibration curves for $NH_4NO_3$ in water (no soil), to verify our soil measurements are from $NH_4^+$ alone. On the chemPEGS we tested 3 concentrations of $H_2SO_4$ functionalization. Higher concentration $H_2SO_4$ takes longer to neutralize by the same amount of $NH_4^+$ . The 10µl 0.025M $H_2SO_4$ functionalization was the best compromise between precision and measurement time.

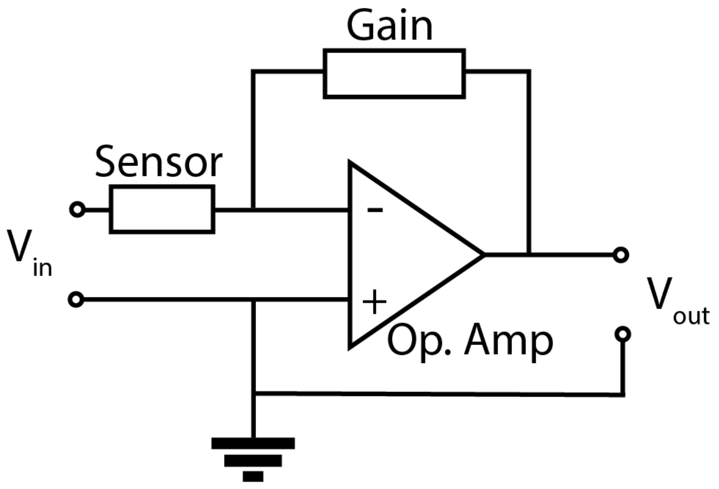


**Figure S2**

Schematic showing home-made electronics for measuring ionic impedance during neutralization on chemPEGS (Sensor). An alternating voltage was supplied ($V_{in}$) across the chemPEGS, and the current passing through was measured as a voltage ($V_{out}$) with a transimpedance amplifier (implemented with an Operational Amplifier), amplified with a resistor (Gain).

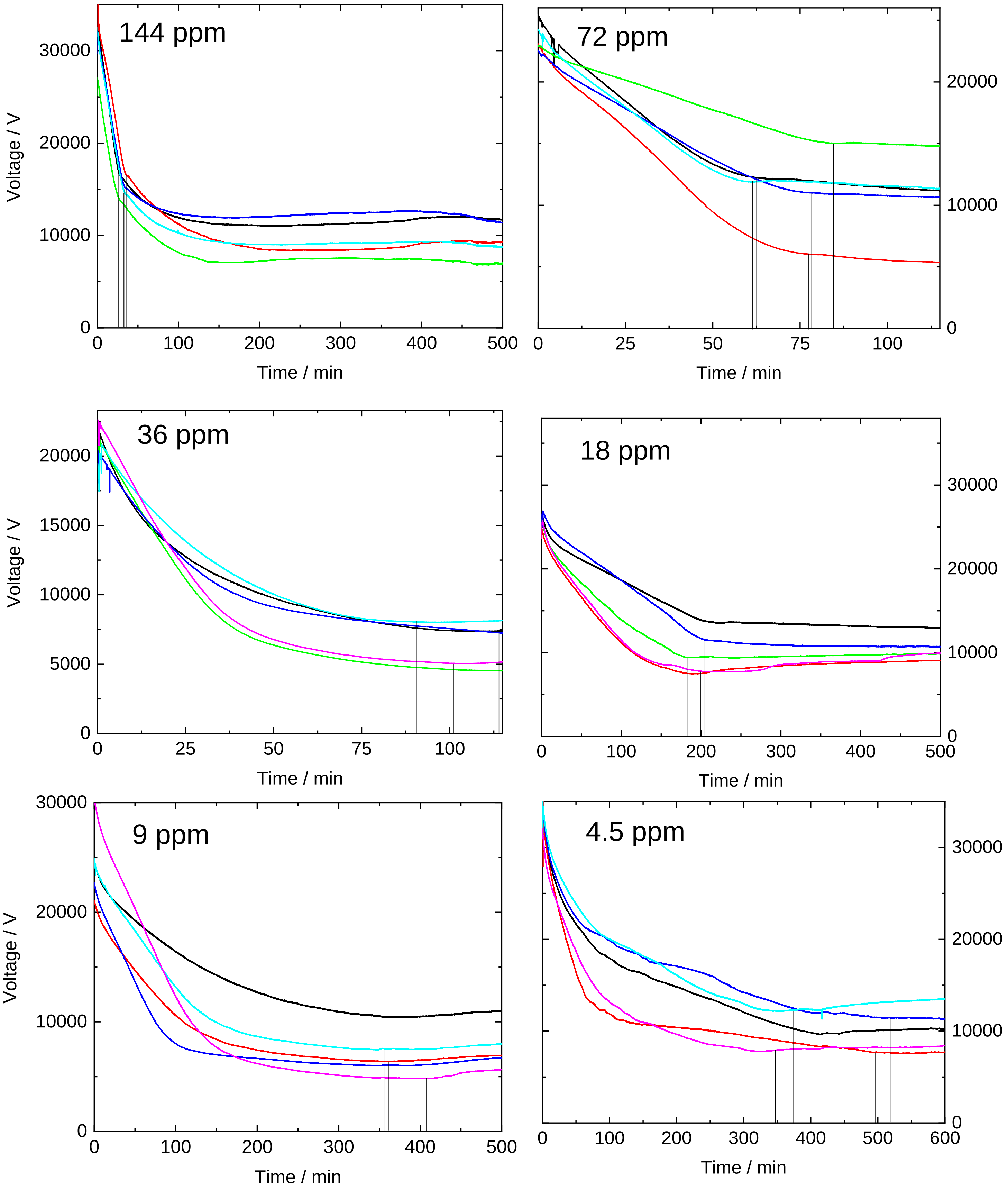
144 ppm
Voltage / V
0
10000
20000
30000
0
100
200
300
400
500
Time / min
72 ppm
0
10000
20000
0
25
50
75
100
Time / min
36 ppm
Voltage / V
0
5000
10000
15000
20000
0
25
50
75
100
Time / min
18 ppm
0
10000
20000
30000
0
100
200
300
400
500
Time / min
9 ppm
Voltage / V
0
10000
20000
30000
0
100
200
300
400
500
Time / min
4.5 ppm
0
10000
20000
30000
0
100
200
300
400
500
600
Time / min

**Figure S3**

Raw data showing decrease in ionic conductivity (measured as a voltage), as $H_2SO_4$ in the paper scrubber is neutralized by $NH_3$ gas. The moment at which the rate of neutralization slows is recorded (marked in vertical black lines on the Time axis). The time used as analytical signal was A) when the gradient was zero or B) when there was a step change in gradient (>80% change in <10 minutes), which ever happened sooner. Higher concentrations (>144ppm) are not adequately separated, but could be with higher concentration of $H_2SO_4$ in the chemPEGS scrubber, however, this would increase measurement time (see **Figure S1**).

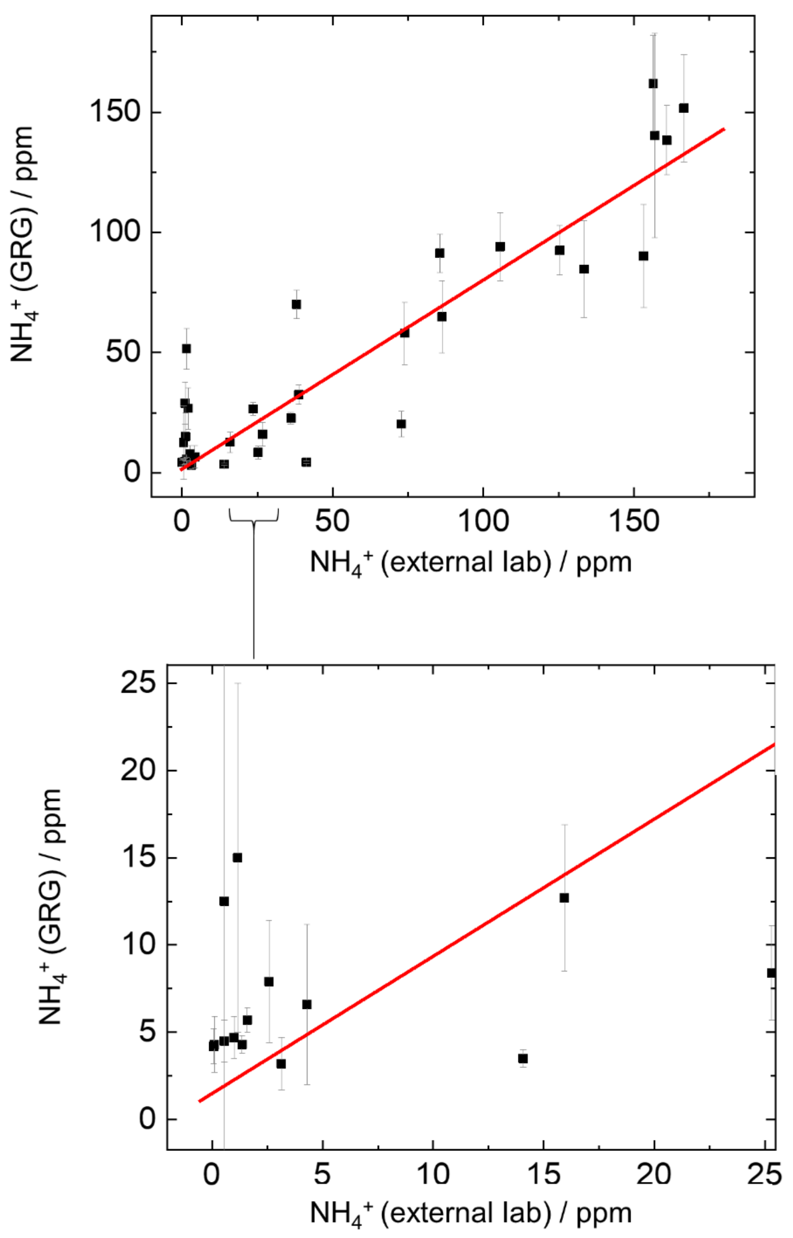


**Figure S4**

After calibration shown in **Figure 2.4** with data from **Figure S3**, 35 soil-$NH_4^+$ measurements were made in soil fertilized with $NH_4NO_3$ for a variety of weather conditions (**see Figure 3**). Soil-$NH_4$+ measurements from our gas-phase $NH_4^+$ sensor (GRG) compared to external laboratory measurements with a score of $R^2$= 0.85.

**Calculation of Fertilization Rate [kg/ha] from Concentration [ppm] and Soil Parameters**

Fertilization Rate[kg/ha] = Hectare Area[$m^2$] × Sample Depth[m] × Density[kg/$m^3$] × Concentration[ppm]

Hectare Area = 10,000 $m^2$

Sample Depth = Pot height = 0.26 m

Soil Density = 774 kg/$m^3$

Concentration of $NH_4NO_3$ = 120 ppm

10,0000 × 0.26 × 774 × (120/1,000,000) = 241 kg/ha

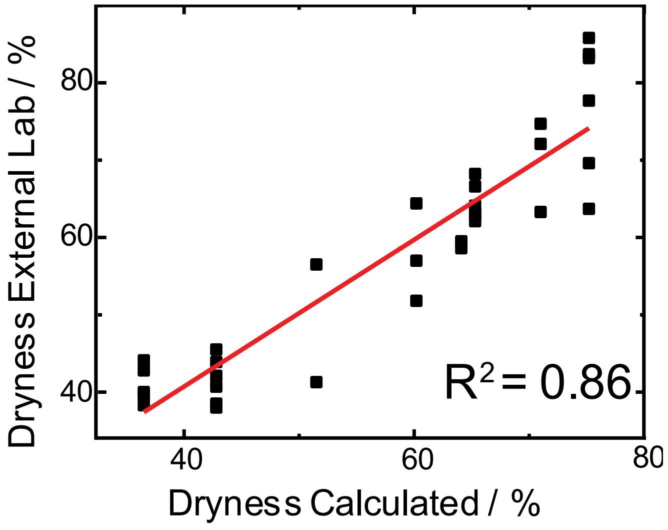


**Figure S5**

In our dataset, dryness is highly correlated with rainfall and temperature. Dryness can be predicted using linear regression with $R^2$ = 0.86 using the equation below, and hence can be estimated using these two metrics without needing further analytical measurements.

$$Dryness[\%] = 0.8853\, Temperature[°\mathrm{C}] - 3.0373\, Rainfall[\mathrm{mm}] + 49.3928$$

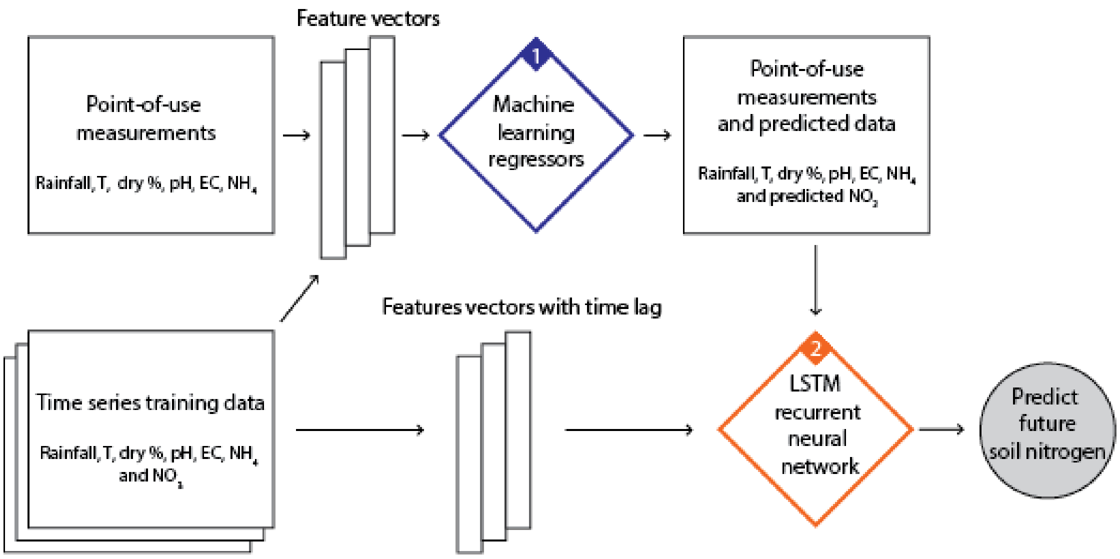


**Figure S6**

Machine learning models trained with readily available environmental data and PoU measurements enable prediction of difficult-to-measure soil nitrogen instantaneously with machine learning regressors (**Figure S6.1**). Prediction into the future is then enabled by treating data as time series in a LSTM neural network (**Figure S6.2**).

**Data Processing for Machine Learning**

The following steps were taken to predict instantaneous soil- $NO_3^-$ (**Figure S6.1**)

1. Anomalous data removed
2. Data for missing days interpolated smoothly with Akima spline
3. Time series length capped at 16 days
4. Data normalized over range 0-1
5. Feature selection: Features ranked for predicting $NO_3^-$ calculated by weight and gain using XGBoost – see **Figure 4.1**. Combinations are tested starting with all 7 features, then only 6 most important features, then the most important 5, down to the single most important feature

6. Cross-validation: Data from each time series is removed from the training data set sequentially, and used as test data instead
7. Regressor selection: Random Forest, Gradient Boosting, Adaboost, Extra Trees, Knn and XGBoost were tested
8. Hyperparameter tuning: Grid search used to tune up to 3 hyperparameters for each regressor

The following steps were taken to predict soil-$NH_4^+$ and soil-$NO_3^-$ 1-12 days into the future (**Figure S6.2**)

1. Concatenate all time series into one multivariate time series
2. Normalize data $-1 < x < 1$ (for a bounded and stable neural network computation)
3. Format time series data for supervised learning (generate additional input features with time lag, generate output, which is soil-$NH_4^+$ and soil-$NO_3^-$ at future time)
4. Remove each time series sequentially and train model on remaining data
5. Predict the removed time series from its measurements on Day 0
6. Create LSTM model and tune hyperparameters with grid search (time lag, epochs, batch size, number of neurons)
7. Record score (mean squared error) of predictions, repeat each prediction 7 times and compare average scores to determine optimal tuning, train LTSM model with optimal tuning to generate predictions of $NH_4^+$ and $NO_3^-$ into the future

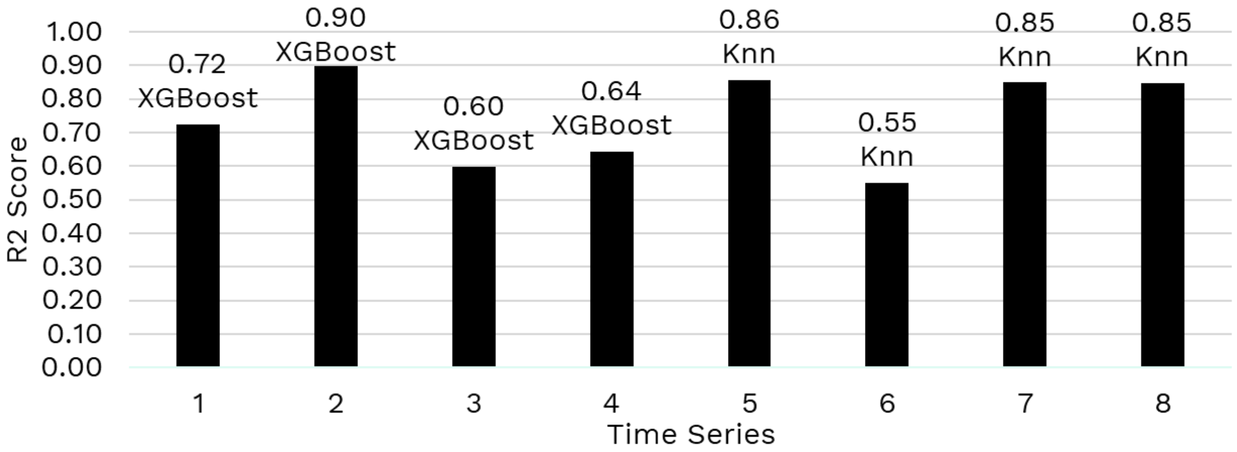


**Figure S7**

Each time series 1-8 (corresponding to soil under different environmental conditions, by controlling rainfall and temperature **-** see **Figure 3**) was sequentially removed from the data, and predicted using the remaining data. Testing all combinations of features, regressors and tuning parameters results in a best-case prediction score for each time series, shown here. XGBoost is gives the best predictions for dryer soils and Knn for wetter soils. Poorer $NO_3^-$ predictions at moderate (3-5mm) rainfalls, where $NO_3^-$ levels were relatively constant over time. The number of features (n_features), regressor and optimal tuning parameters for each time series are listed below.

1. n_features = 5, XGBoost (eta = 0.0, gamma = 0.0, max depth = 6.0)
2. n_features = 3, XGBoost (eta = 0.0, gamma = 0.0, max depth = 6.0)
3. n_features = 4, XGBoost (eta = 0.0, gamma = 0.0, max depth = 7.0)
4. n_features = 7, XGBoost (eta = 0.0, gamma = 0.0, max depth = 1.0)
5. n_features = 4, Knn (n neighbors = 8.0, leaf size = 1.0, p = 1.0)
6. n_features = 7, Knn score (k = 50.0, leaf size = 1.0, p = 3.0)
7. n_features = 7, Knn score (k = 14.0, leaf size = 1.0, p = 1.0)
8. n_features = 6, Knn score (k= 20.0, leaf size = 1.0, p = 10.0)

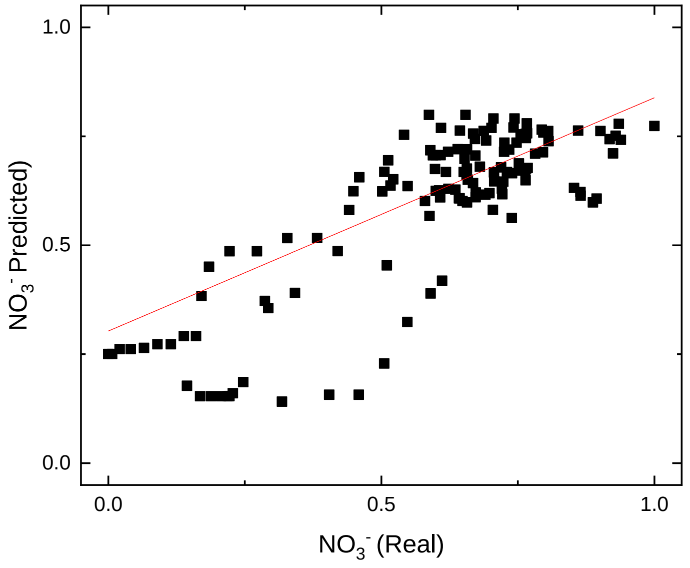


**Figure S8**

The same process was followed as **Figure 4.1** (bottom left), with the same tuning parameters (k=14, leaf size=1, p=1), but test inputs were from the external lab rather than PoU sensors from our lab, thus removing the impact of inaccuracy from our lab. The resulting score is $R^2$=0.68 (compared to the $R^2$=0.63 with our lab inputs).

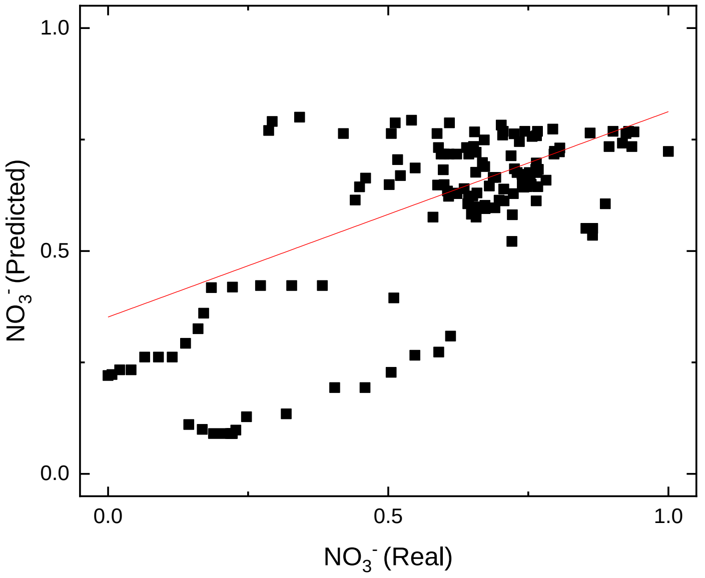

**Figure S9**

Knn predicts $NO_3^-$ using a model tuned (k=11, leaf size=3, p=20) to only the most basic inputs – days since fertilization, rainfall and temperature (i.e. requiring no soil sensors at all.) with $R^2 = 0.54$. One model was used for all environmental conditions.

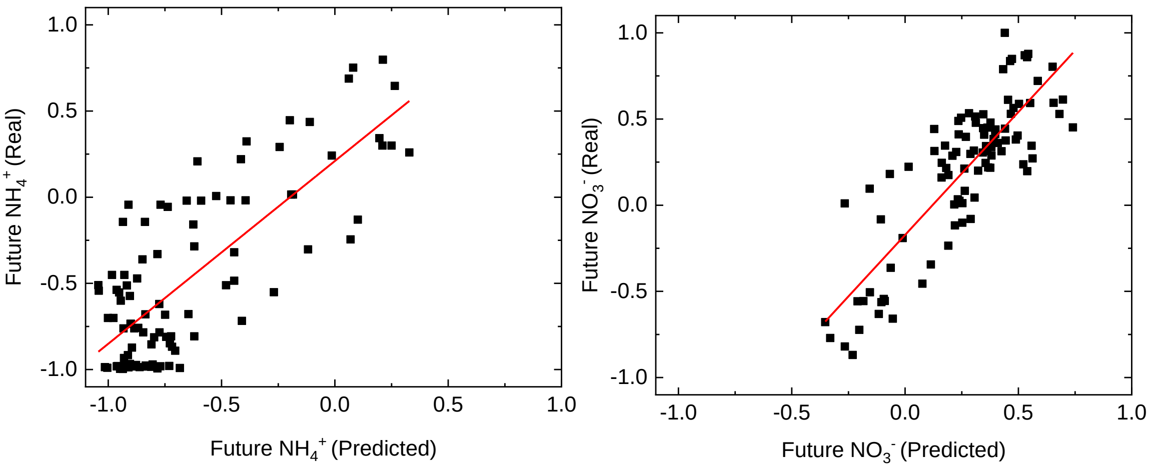


**Figure S10**

$R^2$ score for data in **Figure 4.2**, showing soil-$NH_4^+$ and soil-$NO_3^-$ predicted by LSTM ML model 1-12 days into the future. The LSTM ML model was poorer at predicting lower soil-$NO_3^-$ levels and higher soil-$NH_4^+$ levels.

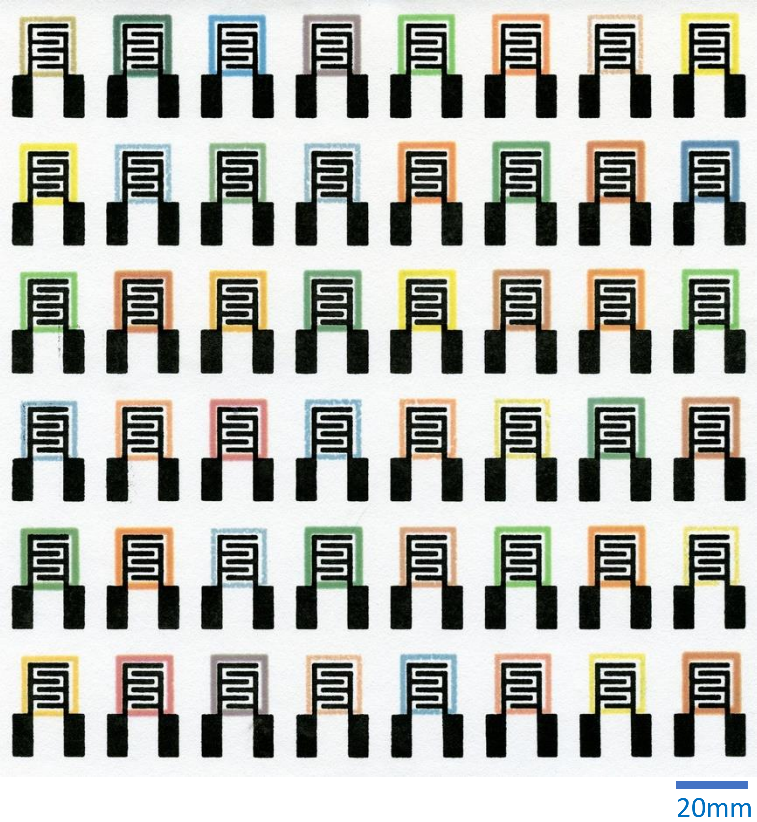


**Figure S11**

Array of batch processed chemPEGS, comprising interdigitated carbon electrodes on chromatography paper, surrounded by a colorful printed wax barrier.
